# Heterogeneity and genomic evolution of metastatic prostate cancer

**DOI:** 10.1101/2023.09.01.556000

**Authors:** Sijia Wu, Zhennan Lu, Yanfei Wang, Xiaobo Zhou, Liyu Huang

**Author notes:** **Correspondence:** Corresponding Author, E-mail addresses (L.H.), (X.Z.). These authors contributed equally to this work (Sijia Wu, Zhennan Lu).

## Abstract

**Background:** Metastasis is the primary cause of prostate cancer-related deaths. However, the underlying molecular mechanisms and evolutionary patterns remain largely uncharacterized.

**Methods:** We evaluate the heterogeneity and genomic evolution of prostate cancer with multi-organ metastases. The samples include 32 primary samples, 23 lymph node metastases, 22 bone metastases, 16 liver metastases, and four pelvic mess metastases. They are analyzed to identify the mutated genes enriched in metastatic samples, selected by metastases, and leading to different long-distance migrations. These metastasis-related alterations constitute a Mscore for evaluating the metastatic risk of primary prostate tumors.

**Results:** Our analysis discovers 21 metastasis-related mutated genes in total. Of them, 14 genes are finally selected for metastatic risk prognosis, including the mutations of *AR* and *KMT2C* with high prediction ability. A Mscore established with these 14 characteristics by the xgboost model displays its ability to classify primary tumors and metastases. This score can further divide primary prostate tumors from the TCGA cohort into two groups. The two subsets present significantly differential survival risks. This score can also identify metastasis-featured primary tumors for breast cancer, bladder cancer, liver cancer, and uterine corpus endometrial carcinoma.

**Conclusion:** Our research proposes 14 molecular features potentially driving prostate cancer metastasis. The Mscore established on them can estimate the metastatic risk of primary tumors.

## 1 Introduction

Prostate cancer is one of the most common cancers in men worldwide [1]. About 20% of men diagnosed with primary prostate cancer will get metastatic cancer during their lifetime [2]. The current treatment for metastatic prostate cancer is androgen deprivation therapy. However, nearly all patients develop resistance to it, a state known as metastatic castration-resistant prostate cancer. The insufficient treatment effectiveness is due to the unclear molecular mechanisms and evolutionary patterns of prostate cancer metastasis [3].

High-throughput technologies such as whole-exome sequencing (WES) enable the identification of molecular signatures related to cancer metastasis [4] by comparing primary and metastatic tumors. For example, androgen receptor (*AR*) activity inversely correlates with tumor cell progression, suggesting mutated *AR* is a metastatic risk of prostate cancer [5]. The alterations identified by this differential analysis present the genomic differences between the primary and metastatic prostate tumors [6]. But they only achieve the area under the receiver operating characteristic curve (ROC) of 0.8 for classifying the two groups when combining all the clinical, histological, and RNA features [6]. More precise metastatic genetic drivers are required for identification with diverse methods. They are useful for understanding the mechanisms of prostate cancer metastasis.

For this goal, three different methods are used in this study to characterize metastatic prostate tumors, considering both interindividual and intraindividual genetic alterations. They are differential analysis between primary and metastatic tumors, identification of clonality changes from primary tumor to metastasis for the same patient, and evolutionary origin analysis using the data of patients with multiple metastases. The identified metastasis-related features are then evaluated by ROC curves of the classification between primary and metastatic samples. Their effectiveness in metastatic risk prognosis of primary tumors is subsequently assessed in additional datasets of prostate and other cancers by survival analysis.

## 2 Materials and methods

### 2.1 Patient samples

There were 72 prostate cancer patients involved in this study (phs000673.v2.p1) [7]. The tumors included 32 primary samples, 23 lymph node metastases, 22 bone metastases, 16 liver metastases, and four pelvic mess metastases, as shown in Figure 1A. They were all analyzed for the heterogeneity of prostate cancer. Of them, seven patients had paired primary and metastatic tumors. The samples from these seven patients constituted 17 different primary-metastasis pairs. These pairs were used in clonality analysis to identify metastasis-selected genetic features. Of the seven patients, three individuals had multiple organ metastases (Table S1). They were applied for the genomic evolution analysis. Additionally, there were also 75 metastatic samples and 11 primary tumors from MET500 (phs000909.v.p1) [8] and 449 primary samples from TCGA [9] involved in this study to validate the discoveries of potential metastatic drivers.

**Figure 1.**
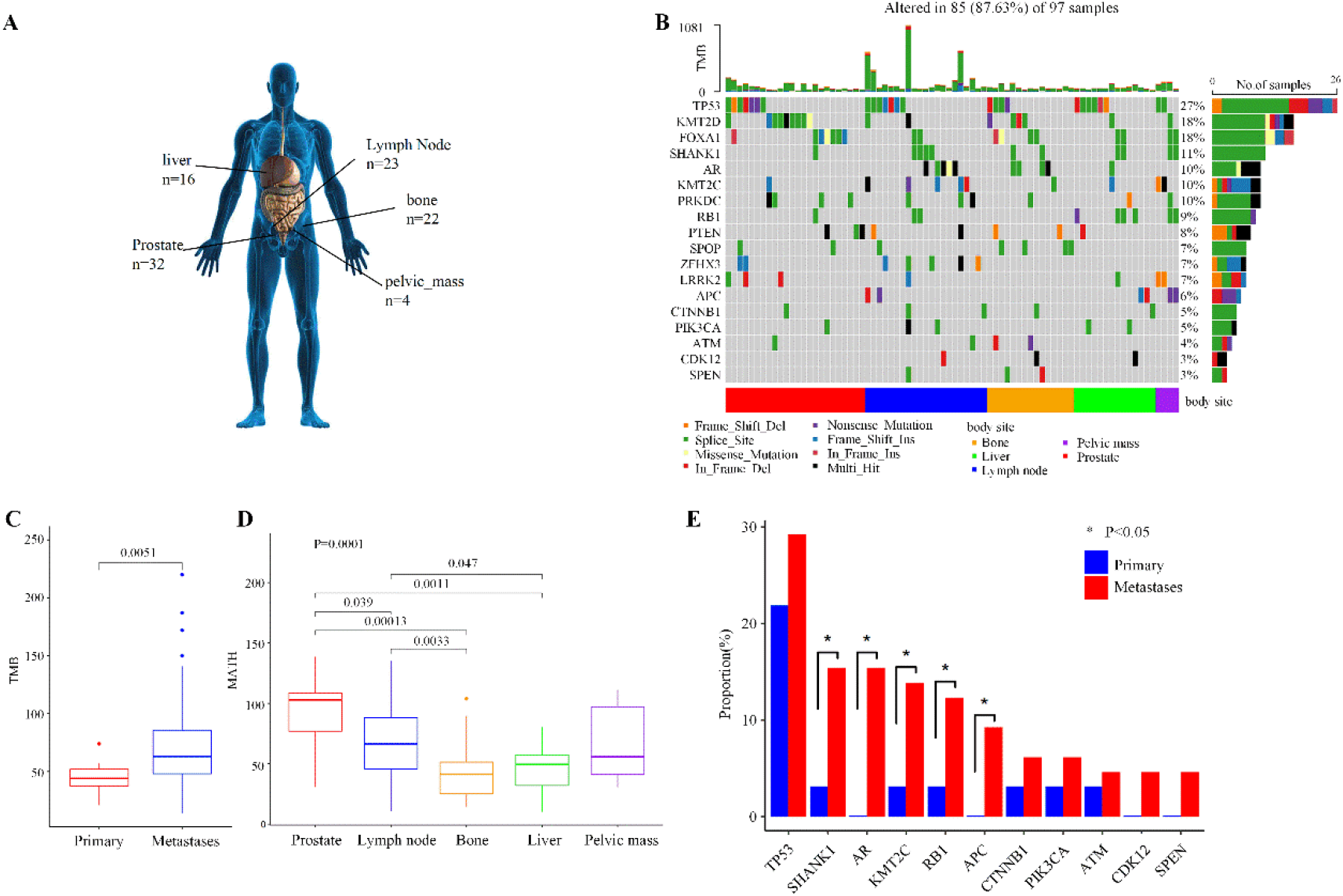
The genetic landscape of metastatic prostate cancers. (A) Schematic diagram showing the biopsy sites of patients and their sample numbers. (B) Genetic profile of all the prostate cancer samples. The heatmap shows the gene mutation landscape for each sample. The top histogram indicates the TMB of each patient. The right bar plot represents the distribution and compositions of mutations for each gene. (C) The comparisons of TMB between primary and metastatic tumors. (D) The comparisons of MATH score among primary prostate tumors and metastases at lymph nodes, bone, liver, and pelvic mass. (E) The comparison of top 11 metastasis-enriched features between primary and metastatic cancers. Wilcoxon tests are used for all the comparisons. TMB, tumor mutational burdens; MATH, Mutant-allele tumor heterogeneity.

### 2.2 Identification of somatic mutations and copy number variants

The whole-exome sequencing data were processed using an in-house pipeline (Figure S1). In detail, FastQC was run on the raw reads to filter out the low-quality samples (q < 25, e > 0.1, and length < 36) [10]. The remaining sequencing reads were aligned to the human reference genome (hg38) with Burrows-Wheeler Aligner (BWA) [11]. These aligned reads were processed by Mutect2 of Genome Analysis Toolkit (GATK, v4.2.5.0) to identify somatic single nucleotide variants (SNV) and small insertions and deletions (indels) [12]. They were also used to detect copy number variations (CNV) by CNVkit (v0.9.9) [13]. The genomic information of these variants was annotated with the ensemble variant effect predictor (VEP, v88.9) [14].

### 2.3 Differential analysis between primary and metastatic prostate tumors

The overall genome differences between primary and metastatic tumors were achieved from their comparisons of tumor mutational burden (TMB) and mutant-allele tumor heterogeneity (MATH) [15]. TMB was the total number of nonsynonymous bases in the 1Mb genomic regions divided by the covered genome using maftools [16]. MATH score was a quantitative assessment of genetic intra-tumor heterogeneity.

Specifically, the differences in each variant between primary and metastatic tumors were evaluated by MutSigCV (v1.41) [17] for somatic mutation and by GISTIC (v2.0.23) for copy number variant [18]. The variants were identified as metastasis-enriched genetic features when fulfilling the significance level of the statistical test (FDR < 0.1) and having mutation frequency greater than 3% and higher in metastatic tumors than primary samples.

### 2.4 Clonalities analysis for primary-metastasis samples from the same patient

The clonalities analysis started from the identification of cancer cell fractions (CCFs) for somatic mutations by PyClone (0.13.1) [19] and the quantification of CCFs for copy number variants by FACETS (v0.6.2) [20]. The variants with CCF greater than 0.6 were clones, and the others were identified as subclones [21]. These clonalities in the paired primary and metastatic samples labelled the variants as one of three groups [22]. The first group contained selected events that were either clonal/subclonal in metastasis and absent from the primary tumor or clonal in metastasis and subclonal in the primary tumor. The second group included maintained events that were either clonal in the primary tumor and clonal/subclonal in metastasis or subclonal in both the primary tumor and metastasis. The third group covered unselected events that were clonal/subclonal in the primary tumor and absent from the metastasis. The last two kinds of events constituted a not-selected group.

The proportions of each variant to be selected or not-selected events in the 17 primary-metastasis pairs were compared by a Binomial test (*P* < 0.05) [22]. Their fractions were then compared between primary and metastatic tumors from the 72 prostate cancer patients and the MET500 dataset using Fisher’s exact test (*P* < 0.05). The variants showing significantly differential results and occurring in more than two tumors (n_i_ ≥ 3) were finally used in the following analysis as metastasis-selected variants.

### 2.5 Evolutionary analysis of metastatic prostate cancer

The three patients with multiple organ metastases (Table S1) were involved in the evolutionary path analysis of metastatic prostate cancer by Treeomics [23] with the maximum likelihood algorithm. The samples grouped into the same branch were determined to have a common origin. The variants potentially driving different distal metastasis were identified as metastasis evolution-associated variants.

### 2.6 Machine-learning model to predict the metastatic risk of primary tumors

With the potential metastatic drivers identified above, an xgboost [24] model was developed to classify primary and metastatic tumors. Its effectiveness was evaluated in the tumors of 72 prostate cancer patients and then validated in the independent MET500 dataset. The significantly metastatic drivers during the training process were integrated to define a Mscore index. It can divide the primary tumors into two subsets showing different metastatic risks, Metastasis-Featuring Primary (MFP) tumors (Mscore > 0.5) and Conventional Primary (CP) tumors (Mscore < 0.5). Its usefulness in predicting metastatic risk was tested in a primary prostate tumor cohort and other primary cancers from TCGA with survival analysis and aggressiveness evaluation by Gleason score.

## 3 Results

### 3.1 The heterogeneity between primary and metastatic prostate tumors

Our pipeline identified 248 somatic single nucleotide variants (SNVs) and 163 somatic copy number variants (CNVs) across the tumors of 72 prostate cancer patients. Of the genes with somatic SNVs, the most frequently mutated ones were *TP53* (27%), *KMT2D* (18%), *FOXA1* (19%), *SHANK1* (11%), *AR* (10%), and *KMT2C* (10%) (Figure 1B). Of the somatic CNVs in Figure S2, the most frequent CNVs included losses at 17q21.31, 16q22.1, and 6q15 regions and gains at 8q24.3 and 12q13.2 regions. These results were consistent with the analyses conducted in previous studies (Figure S3) [25-28]. The analyses confirmed the reliability of our pipeline.

The analysis of these variants revealed that the metastatic samples had high tumor mutational burden (TMB, Figure 1C) and low tumor heterogeneity (MATH, Figure 1D). It showed newly occurred and similar mutations when tumors metastasized to liver, bone, and other organs. The comparisons of the variants between primary tumors and metastases also provided 11 metastasis-enriched features (Figure 1E). They included mutated *TP53, SHANK1, AR, KMT2C, RB1, APC, CTNNB1, PIK3CA, ATM, CDK12*, and *SPEN*. Specifically, *AR* mutations were rare in primary prostate tumors but common in metastatic prostate cancer. *KMT2C* mutations were significantly high in metastatic samples. Combined with previous studies [26, 29], these results suggested the 11 features as potential metastatic drivers. All the above results revealed the genomic heterogeneity between primary and metastatic prostate tumors.

Moreover, genetic differences existed not only between primary tumors and metastases but also among diverse distant sites of metastases from prostate tumors. Specifically, the MATH value of lymph node metastases was the highest compared to other metastatic sites (Figure 1D). This observation may indicate that a portion of the metastases involved lymph nodes as nodes of dissemination. And the lymph node metastases had more variants (Figure S4), suggesting a higher degree of genomic instability and molecular alterations contributing to the metastatic process through lymph nodes. Additionally, the gene of *AR* was frequently mutated in lymph node metastases (22%, Figure S4B) and bone metastases (18%, Figure S4C). It may describe the potential roles of *AR* mutations in the progression of bone and lymph node metastases. These results highlighted the diverse mutational profiles and genetic heterogeneity across different metastatic sites.

### 3.2 Selected features potentially promoting metastatic progression

The metastasis-selected features were events occurring newly in metastases or changing from subclonal to clonal when primary tumors migrated to metastases (Figure 2A). The statistical analysis of the fractions between the selected and not-selected events for all the features identified 21 genetic alterations in total (Figure 2B). Of these variants, 11 showed significant differences between primary and metastatic samples of the 72 prostate cancer patients (Figure 2C). Six alterations of them had validated their potential functions in another MET500 cohort (Figure 2D). They were the variants of *OBSCN, KMT2C, NR2E1, CPD*, 6q15 loss, and 8q24 gain.

**Figure 2.**
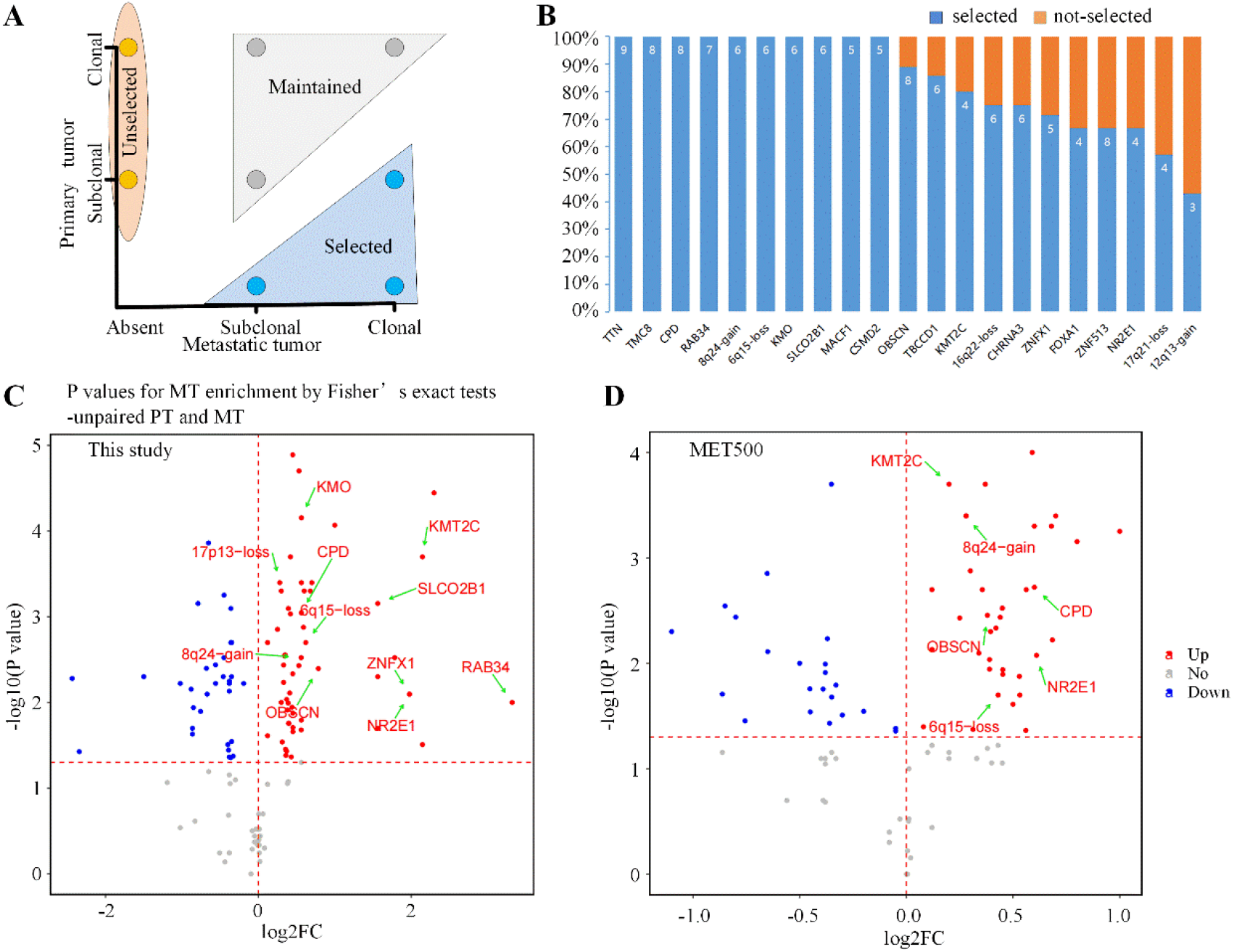
Identification of metastasis-selected features. (A) Schematic illustration for the definition of selected, maintained, and unselected events in metastases. (B) The proportions of selected events for each variant. These variants occur in more than two tumors and have differential fractions of selected events compared to not-selected events in the 17 primary-metastasis pairs. (C) The comparison of variants between primary and metastatic samples of the 72 prostate cancer patients by Fisher’s exact test. The x-axis represents the log values of the mutation ratio in metastases to that in primary samples. (D) The comparison of variants between primary and metastatic samples from the MET500 dataset. PT, primary tumor; MT, metastatic tumor; FC, fold change.

Then the analysis of these metastasis-selected molecular features between primary tumors and different metastases also revealed the metastasis site-specific variants (Figure S5). For example, the *KMT2C* mutations may drive lymph node metastases. The metastasis-selected molecular features may not only drive prostate cancer metastasis but also contain information about the metastatic potency to different organs.

### 3.3 Two evolutionary patterns of prostate cancer metastases

The above MATH analysis indicated the possible involvement of lymph nodes as nodes of the metastasis pathway (Figure 3A). The phylogenic trees of C0 and C89 patients also re-addressed this hypothesis (Figure 3B-C). They showed the clusters of distant metastatic sites and lymph node metastases in the same phylogenic clade. It revealed the seeding of the distal metastases through earlier colonized lymph node metastases. All the analyses described that cancer cells spread across lymph nodes to the liver, bone, and other organs.

**Figure 3.**
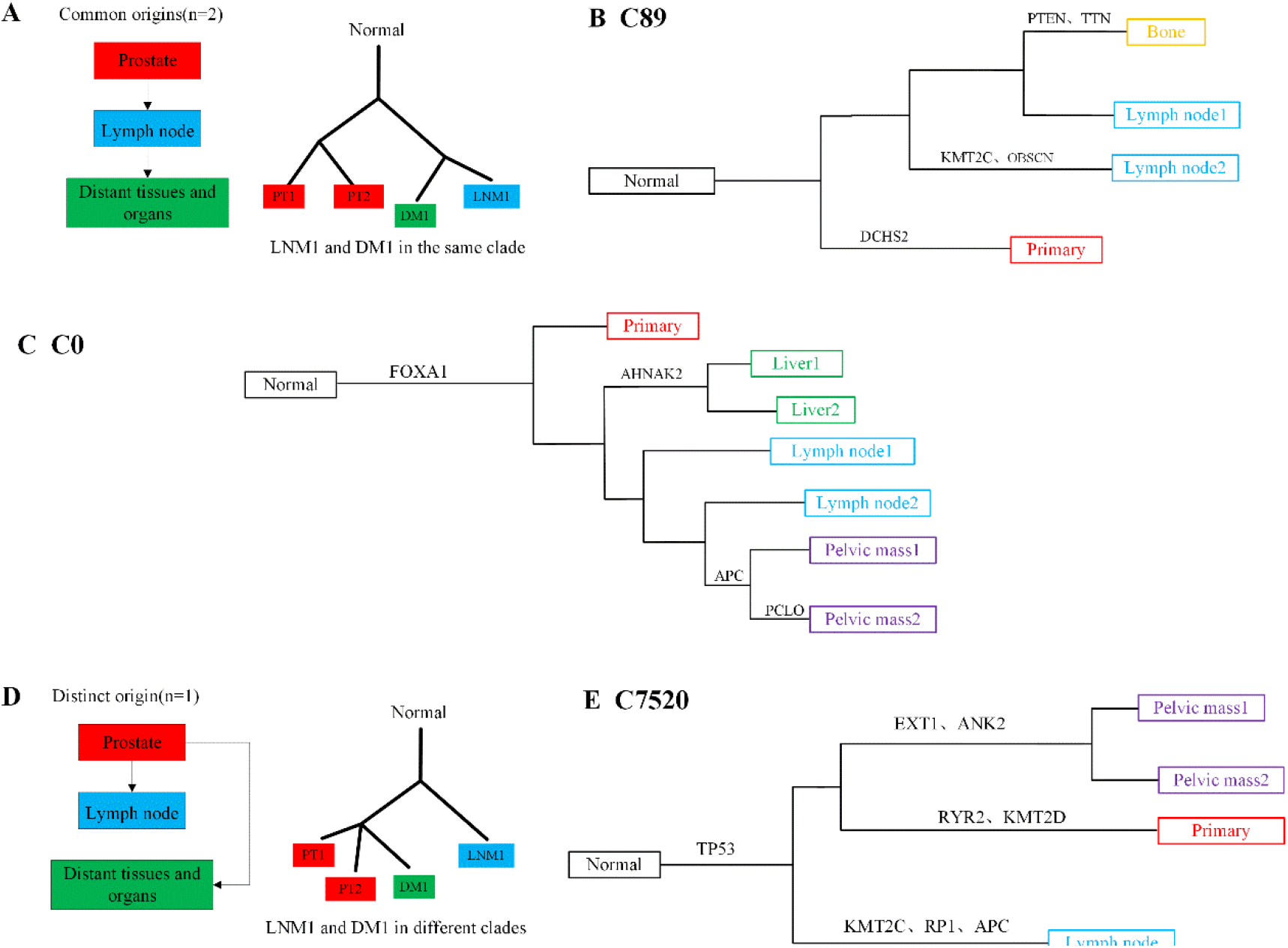
Evolutionary analysis of prostate cancer metastases. (A) One possible tumor migration route (left panel) and phylogenic relationship (right panel) of prostate cancer. (B) The phylogenic tree of C89 with canonical prostate cancer drivers. (C) The phylogenic tree of C0 with canonical prostate cancer drivers. (D) Another possible tumor migration route (left panel) and phylogenic relationship (right panel) of prostate cancer. (E) The phylogenic tree of patient C7520 with canonical prostate cancer drivers.

Additionally, the distant metastases of the C7520 patient tended to have a common evolutionary origin with primary tumors rather than through the lymph node metastases (Figure 3D-E). It revealed an additional pathway that metastases in different organs possibly originated from primary tumors independently.

The evolution analysis identified ten genes as metastasis evolution-associated mutated genes (Figure 3). Specifically, *KMT2C* was a mutated gene specific to lymph node metastasis for C89 and C7520 patients. It was consistent with the above analysis showing a higher proportion of *KMT2C* mutations in lymph node metastasis (Figure S5). This variant was recognized as a potential metastatic driver with all the above three methods. It required further deep analysis and discussion.

### 3.4 Mscore for metastatic risk prognosis of primary prostate cancers

The above analyses totally discovered 21 genes potentially driving prostate cancer metastasis. They can accurately identify the metastatic samples from primary tumors with an xgboost model (Figure 4A) in the datasets of 72 prostate cancer patients (AUC = 0.865, Figure 4B) and MET500 (AUC = 0.845, Figure 4C). The high classification results revealed the relevance of these genetic alterations to metastasis. Their contributions to primary-metastasis classification finally determined 14 mutated genes with metastatic promotion evidence (Figure 4D). *TP53* mutation was suggested here as a metastatic feature with high prognosis ability, followed by the well-known mutations of *AR* [7], *RB1* [30], and *KMT2C* [26], and *OBSCN* [31] (Figure 5D).

**Figure 4.**
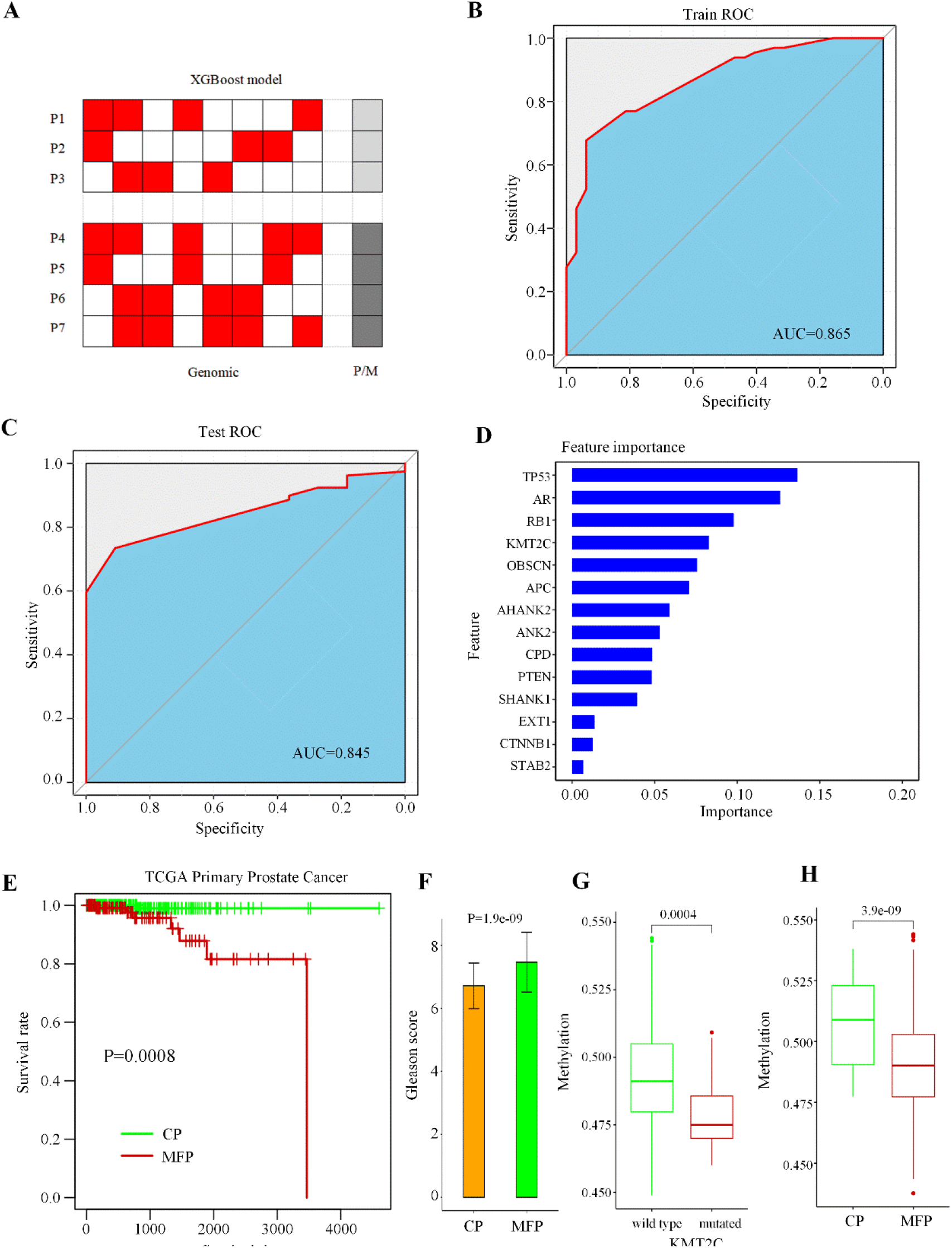
Identification and evaluation of metastasis-featuring primary tumors. (A) An XGBoost model with different genomic features and the labels of P or M for each sample. (B) The ROC and AUC for the model on the samples of the 72 prostate cancer patients. (C) The ROC and AUC for the model on the samples from the MET500 dataset. (D) The contributions of the 14 genetic alterations to the classification between P and M. (E) KM plot displays the difference in disease-free survival between MFP and CP primary prostate tumors from TCGA. (F) The comparison of Gleason scores between CP and MFP categories. (G) The comparison of methylation levels between primary prostate tumors with mutated *KMT2C* and the wild types. (H) The comparison of the methylation levels between CP and MFP primary prostate tumors from TCGA. P, primary tumor; M, metastatic sample; ROC, receiver operating curve; AUC, area under the curve; CP, conventional primary; MFP, metastasis-featuring primary.

**Figure 5.**
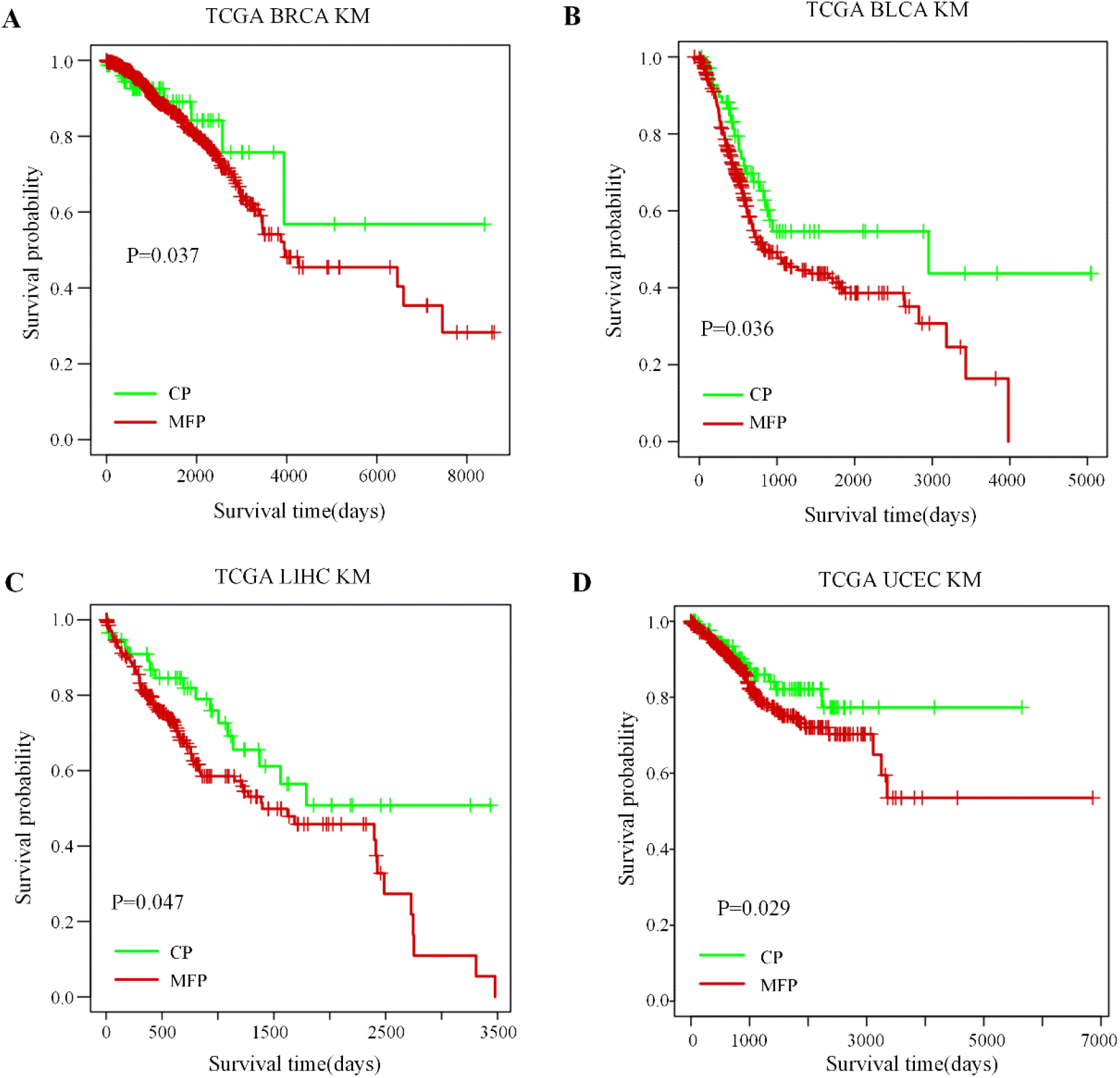
The performance of the Mscore in the other four cancer types from TCGA. (A) The difference in disease-free survival between MFP and CP categories of BRCA primary cancer. (B) The difference in disease-free survival between MFP and CP categories of BLCA primary cancer. (C) The difference in disease-free survival between MFP and CP categories of LIHC primary cancer. (D) The difference in disease-free survival between MFP and CP categories of UCEC primary cancer. BRCA, breast cancer; BLCA, bladder urothelial carcinomas; LIHC, hepatocellular carcinomas; UCEC, endometrial carcinomas.

These 14 mutated genes were integrated as a Mscore index. It was able to estimate the likelihood of one primary tumor to be metastatic. The primary tumors with a Mscore larger than 0.5 were Metastasis-Featuring Primary tumors (MFP) with high metastatic risk. The Mscore was then validated in 449 primary prostate tumors from TCGA. The MFP patients had a poorer survival prognosis of disease-free (*P* = 0.0008, log-rank test, Figure 4E) and higher aggressiveness characterized by mean Gleason score (*P* < 0.001, Figure 4F). All the results demonstrated that the Mscore covered informative metastatic features of prostate cancer.

Specifically, of the 14 mutated genes, *KMT2C*, encoding histone methyltransferases, showed its deficiency to cause DNA methylation in metastasis [32]. A reduced methylation level was also observed in the prostate cancer patients with *KMT2C* mutations (Figure 4G) and the MFP patients (Figure 4H) from the TCGA database. The prostate cancer patients with *KMT2C* mutations had poorer survival risks (Figure S6A). The analyses validated the intrinsic function of this gene and the pro-metastatic role of its deficiency in prostate cancer through the downregulation of methylation.

### 3.5 Mscore for metastatic risk prognosis of other primary cancers

The Mscore also presented its effectiveness in metastatic risk prognosis for breast cancer, bladder urothelial carcinomas, hepatocellular carcinomas, and endometrial carcinomas. The primary tumor patients of these cancers had a higher risk of disease-free survival when they got a high Mscore index (Figure 5). It suggested a generative capacity of the Mscore to stratify cancer patients showing differentially metastatic risks. Of the metastatic features covered in this Mscore, the alterations of *KMT2C, TP53*, and *AR* showed strong prognosis ability. Their positive associations with the Mscore were observed in all four cancer types (Figure 6A-C). Specifically, the deficiency of *KMT2C* caused hypomethylation in hepatocellular carcinomas (Figure 6D). This aberrant methylation then contributed to hepatic carcinogenesis [33] and metastasis (Figure 6E).

**Figure 6.**
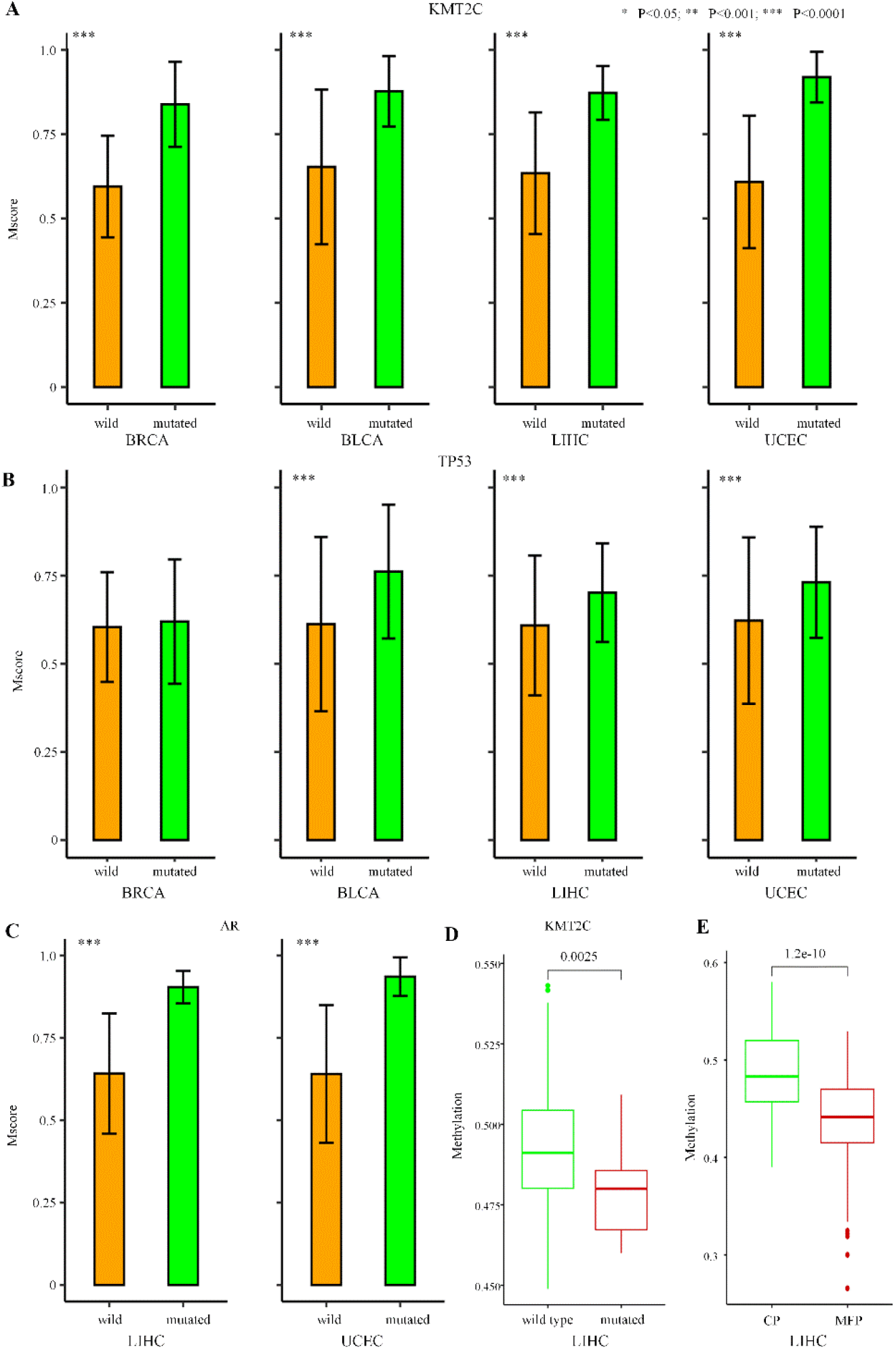
The potential function of *KMT2C, TP53*, and *AR* alterations with metastasis. (A) The comparison of the Mscore between primary tumors with mutated *KMT2C* and the wild types in the four cancer types from TCGA. (B) The comparison of the Mscore between primary tumors with mutated *TP53* and the wild types in the four cancer types from TCGA. (C) The comparison of the Mscore between primary tumors with mutated *AR* and the wild types in the two cancer types from TCGA. (D) The comparison of methylation levels between primary tumors with mutated *KMT2C* and the wild types in LIHC cancer type. (E) The comparison of the methylation levels between CP and MFP primary liver tumors from TCGA.

## 4 Discussion

Primary prostate cancer is a molecularly heterogeneous disease [34-37]. Understanding its metastatic trajectory helps the diagnosis and treatment of this cancer [38]. In this study, the primary and metastatic lesions have various levels of genomic heterogeneity. The analyses identify 14 genetic alterations potentially driving prostate cancer metastases. They present good classification ability between primary prostate tumors and metastatic samples. They can also stratify primary tumors with different metastatic risks for prostate cancer and multiple other cancer types. These variants are useful for personalized therapy.

Of these genetic alterations, the mutation of *KMT2C* is identified as a potential metastatic driver using all three methods. It is a lymph node metastasis-specific genetic alteration and reveals the involvement of lymph nodes in the migration from primary tumors to distant organs. This mutation is associated with metastasis through the downregulation of methylation (Figure 4G-H, Figure 6D-E). It contributes to poor survival prognosis for three cancer types (Figure S6A-C). Moreover, previous studies also report the crucial role of the frequently aberrant *KMT2C* in the occurrence and development of human cancers [39-41]. The pro-metastatic variant can also affect the treatment effectiveness. For example, patients with *KMT2C* mutations [42] receive a shorter treatment time of first-line androgen receptor signaling inhibitors (ARSI, Figure S6D). It means the fast occurrence of drug resistance for these *KMT2C* mutated patients. The detection of *KMT2C* may help clinical decision-making and personalized treatment strategies for cancer patients in the future.

The mutation of *AR* is also another potential metastatic driver. The activation of *AR* alterations occurs only in metastasis (Figure 1E) and contributes to metastatic prognosis (Figure 4D). A higher frequency of *AR* alterations is discovered in patients [26, 42] postexposure to ARSI treatment or taxane chemotherapy (Figure S7A-C). The patients with *AR* mutations [42] receive a shorter ARSI treatment time (Figure S7D). All the above analyses reveal a close association of mutated *AR* with tumor metastasis and drug resistance. It is consistent with previous studies about the enrichment of *AR* mutations in metastatic samples [7, 26, 29, 43, 44] and the promotion role of *AR* deficiency in hormonal therapy escape [45].

During the analysis, several genetic alterations specific to diverse metastatic organs are discovered (Figure S5). Further research on the prediction of metastatic sites is on the plan once got sufficient data. The ratio of the two evolutionary patterns (Figure 3) for prostate cancer metastasis is another further project based on enough patients with multiple organ metastases. These plans, combined with this study here, will help the deep understanding of prostate cancer metastasis and the future design of clinical cancer therapy.

## 5 Declaration

### 5.1 Author Contributions

Conceptualization, S.W.; methodology, S.W., Z.L.; formal analysis, Z.L.; investigation, S.W., Z.L.; data curation, Z.L., Y.W; writing—original draft preparation, Z.L.; writing—review and editing, S.W.; supervision, X.Z. and L.H.; funding acquisition, S.W. and L.H. All authors have read and agreed to the published version of the manuscript.

### 5.2 Informed Consent Statement

The data was obtained from public resources of dbgap (phs000673.v2.p1), MET500 (phs000909.v.p1), and The Cancer Genome Atlas (TCGA).

### 5.3 Data Availability Statement

The data presented in this study are available in the article and Supplementary Material here.

## 5.4 Acknowledgement

The results <published or shown> here are in whole or part based upon data generated by the TCGA Research Network: https://www.cancer.gov/tcga. This research was funded by the National Natural Science Foundation of China (Grant No. 62002270), the Fundamental Research Funds for the Central Universities, the National Natural Science Foundation of China (Grant No. 82227802), the Natural Science Foundation of Shaanxi Province of China (Grant No. 2020JQ-332), the China Postdoctoral Science Foundation (Grant No. 2018M643583), and the National Key R&D Program of China (Grant No. 2017YFA0205202) and partially funded by the National Natural Science Foundation of China (Grant No. 61672422).

