## Supplementary Figure 1-7 and Supplementary Table 1 for "Heterogeneity and genomic evolution of metastatic prostate cancer"

### Supplementary Material

#### 1 Supplementary Figures

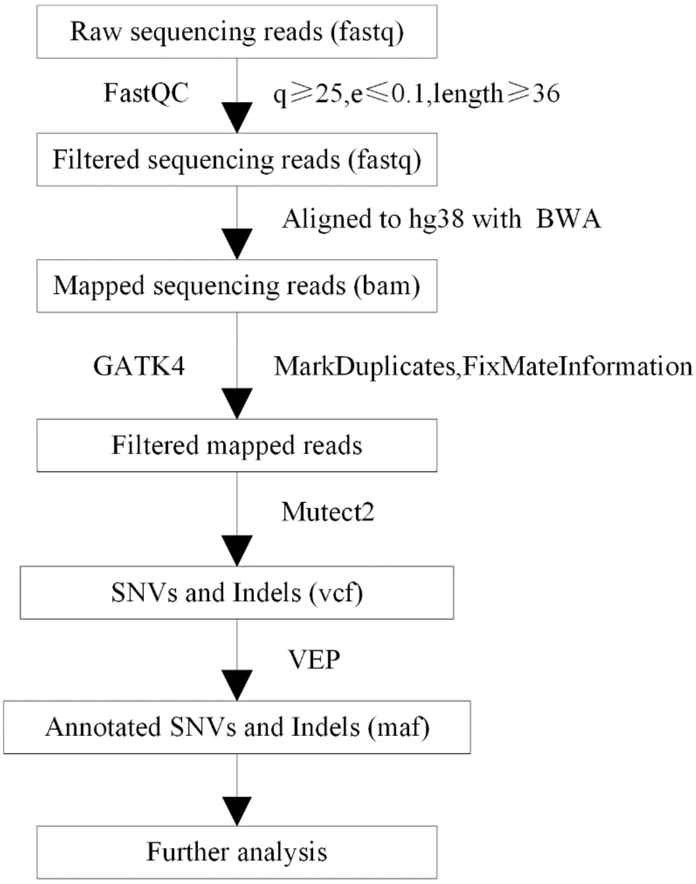


**Figure S1.** Pipeline of somatic mutation analysis. q, read quality; e, error rate; length, read length.

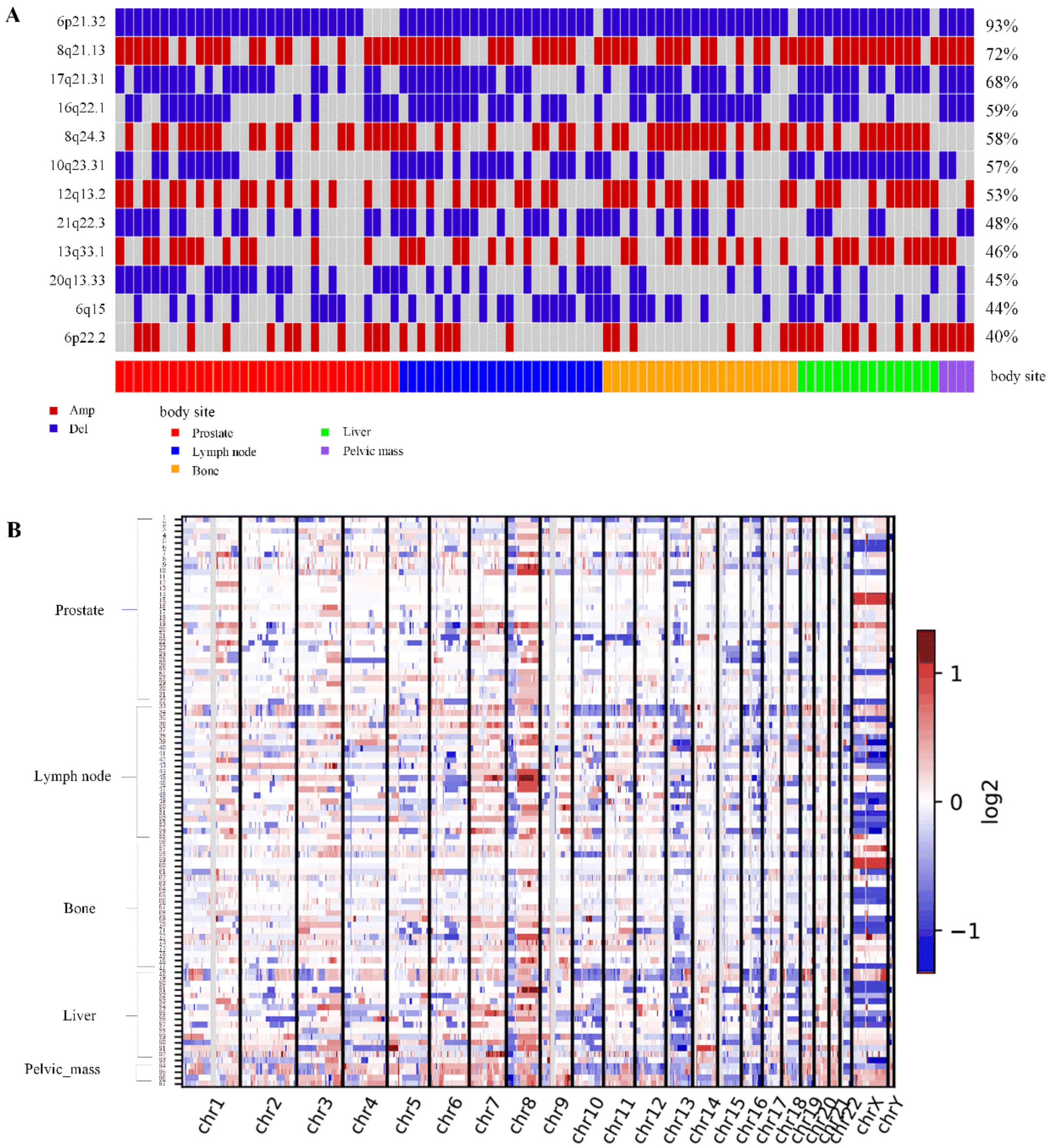


**Figure S2.** The landscape of copy number alterations. (A) The most frequent somatic copy number alterations for the samples of 72 prostate cancer patients. (B) The overview of the copy number alterations for the samples of 72 prostate cancer patients. The red box represents the gain of copy numbers, and the blue box represents the loss of copy numbers.

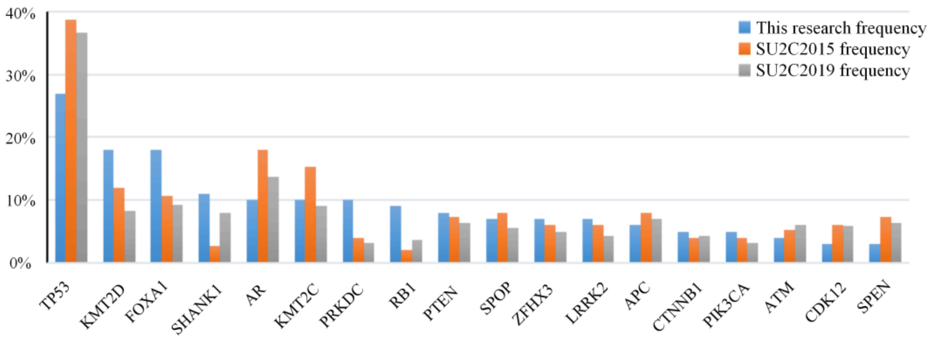


**Figure S3.** The comparison of the top mutation frequencies in different prostate cancer datasets.

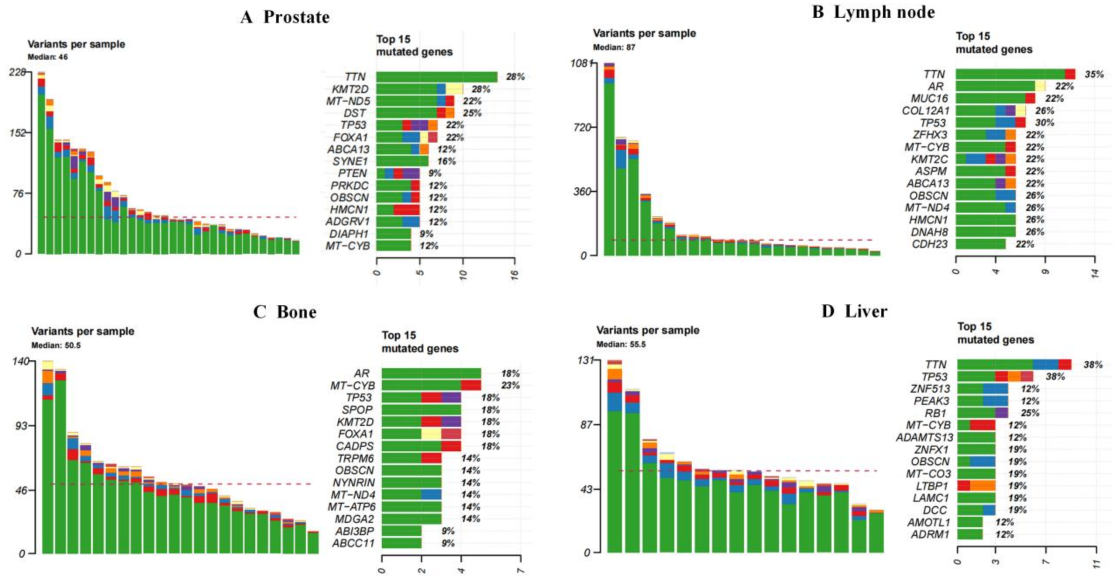


**Figure S4.** The landscape of gene alterations in the datasets of primary and different metastatic prostate cancers. (A) The number of variants per sample (left panel) and the top 15 mutated genes in primary prostate tumors. (B) The number of variants per sample (left panel) and the top 15 mutated genes in lymph node metastatic samples. (C) The number of variants per sample (left panel) and the top 15 mutated genes in bone metastatic samples. (D) The number of variants per sample (left panel) and the top 15 mutated genes in liver metastatic samples. Different colors represent the compositions of the mutations, as shown in Figure 1.

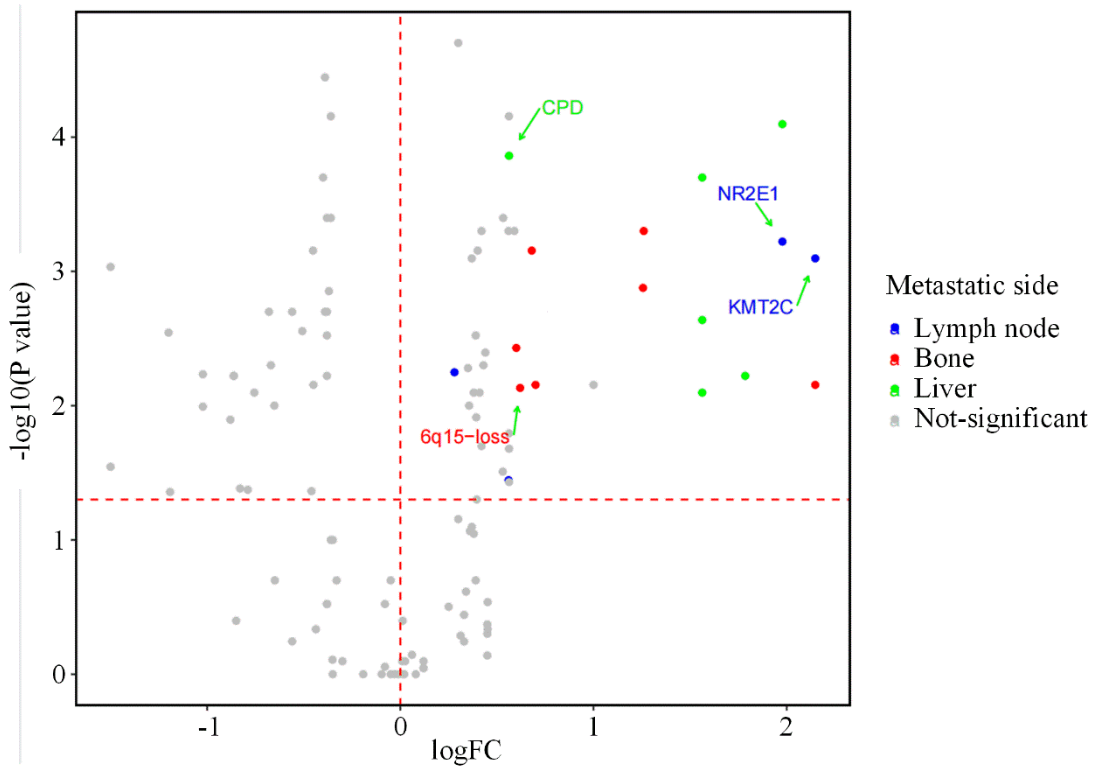


**Figure S5.** The comparison of variants between primary tumors and different metastatic samples by Fisher's exact test.

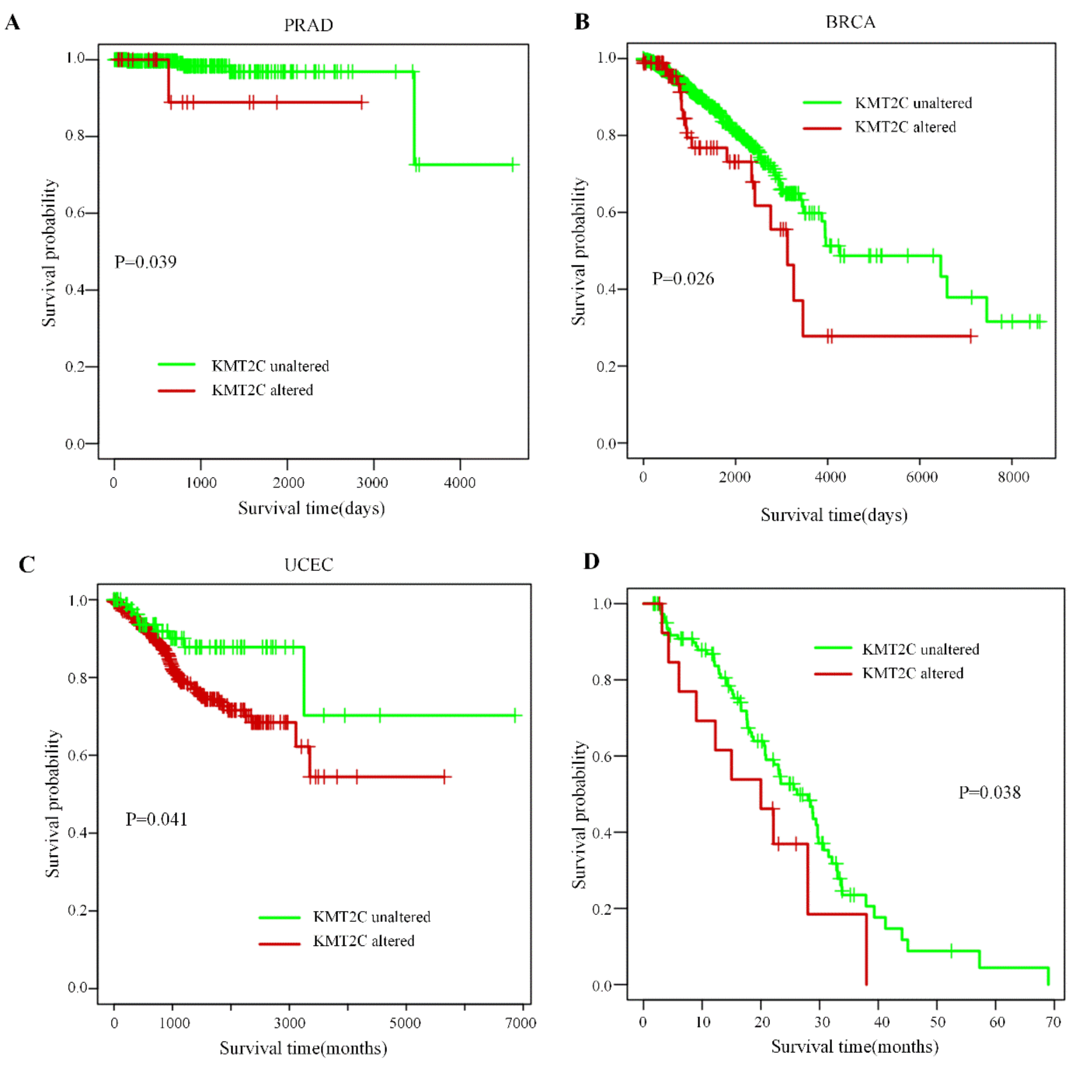


**Figure S6.** Analysis of survival and treatment effectiveness for cancer patients with *KMT2C* alterations. (A) Survival analysis of prostate cancer patients with *KMT2C* alterations. (B) Survival analysis of breast cancer patients with *KMT2C* alterations. (C) Survival analysis of uterine corpus endometrial carcinoma patients with *KMT2C* alterations. (D) The comparison of the first-line ARSI treatment period between patients with *KMT2C* alterations and the wild types. ARSI, first-line androgen receptor signaling inhibitors.

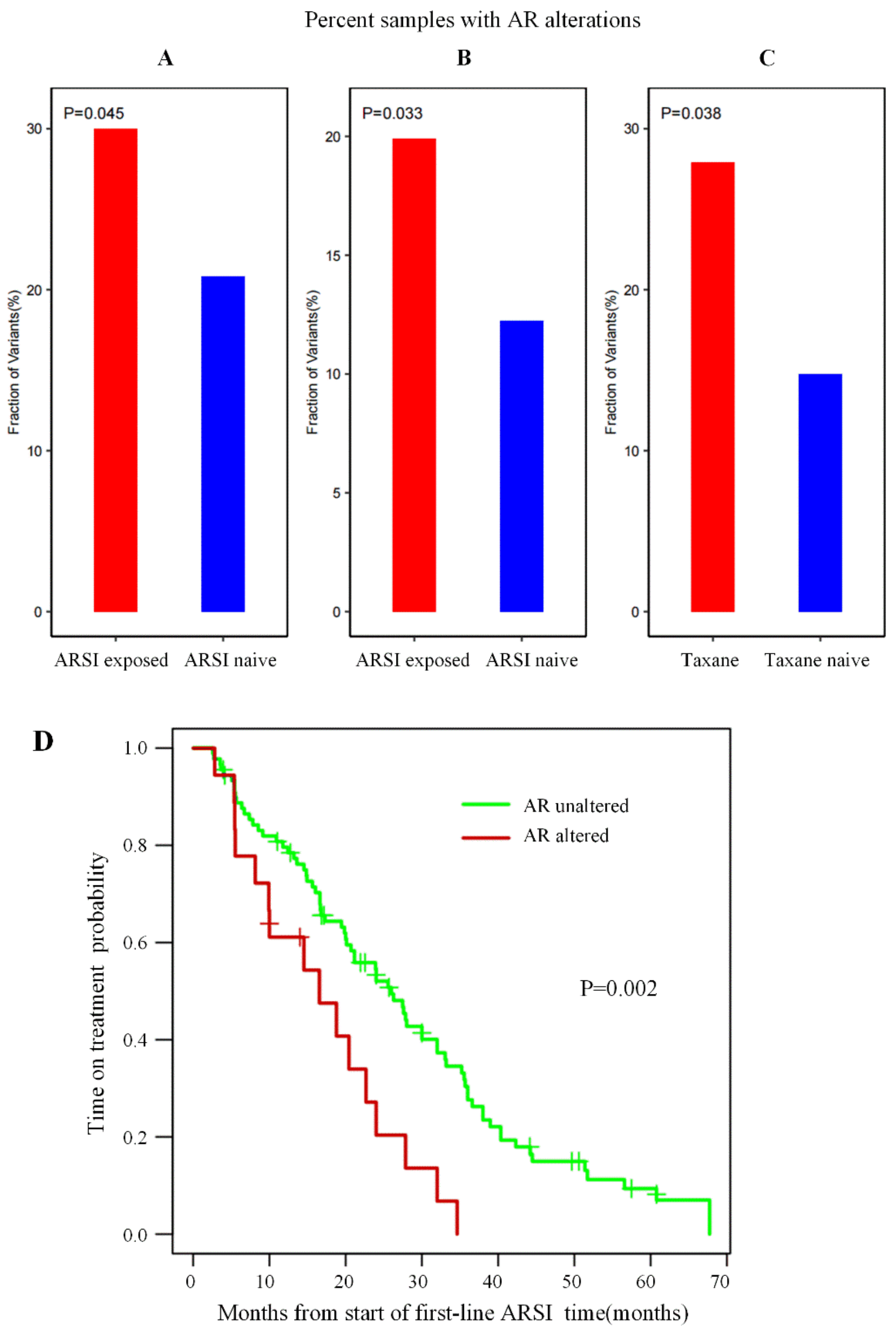


**Figure S7.** The effect of *AR* alterations on drug resistance. (A) The comparison of *AR* alterations between patients exposed to ARSI and ARSI-naive tumors in the dataset of SU2C 2015 cohorts. (B) The comparison of *AR* alterations between patients exposed to ARSI and ARSI-naive tumors in the dataset of SU2C 2019 cohorts. (C) The comparison of *AR* alterations between patients exposed to taxane chemotherapy and taxane-naive tumors in SU2C 2015 cohorts. (D) The comparison of the first-line ARSI treatment period between patients with *AR* alterations and the wild types.

#### 2 Supplementary Tables

**Table S1**. Patients with multiple organ metastases

| Patient | sample | body_site |
|---|---|---|
| WCMC0 | WCMC0_9_LN | Lymph node |
| | WCMC0_6_Li | Liver |
| | WCMC0_10_LN | Lymph node |
| | WCMC0_8_PM | Pelvic mass |
| | WCMC0_1_PM | Pelvic mass |
| | WCMC0_5_Li | Liver |
| | WCMC0_7_P | Prostate |
| WCMC7520 | WCMC7520_3_PM | Pelvic mass |
| | WCMC7520_4_PM | Pelvic mass |
| | WCMC7520_1_LN | Lymph node |
| | WCMC7520_2_P | Prostate |
| WCMC89 | WCMC89_LN1 | Lymph node |
| | WCMC89_P1 | Prostate |
| | WCMC89_LN2 | Lymph node |
| | WCMC89_B | Bone |
